# A Co-culture Cell-Based Reporter Assay for Quantitative Measurement of Integrin αvβ8–Mediated Activation of Latent TGF-β1

**DOI:** 10.64898/2026.06.29.735300

**Authors:** Jianhuan Zhang, Minh Thai, Matthieu Masureel, Cecilia Chiu, WeiYu Lin, Tulika Tyagi, Alessandra Castiglioni, Dhaya Seshasayee, Kelly Loyet

## Abstract

Integrin αvβ8 is a major activator of latent transforming growth factor-β (TGF-β) and an emerging therapeutic target in cancer and immune regulation. However, functional assays that directly measure αvβ8-mediated activation of latent TGF-β in a physiologically relevant context remain limited. Here, we report a co-culture cell-based reporter assay for quantitative measurement of αvβ8-mediated activation of latent TGF-β1. NIH/3T3 reporter cells were engineered to express a SMAD-responsive NanoLuc reporter, constitutive firefly luciferase for internal normalization, and cell-surface GARP–latent TGF-β1. When co-cultured with αvβ8-expressing LN-229 cells, reporter cells produced a robust signal that directly reflected localized latent TGF-β1 activation. The assay demonstrated stable expression of the required biological components, reproducible signal-to-background performance, and sensitivity to benchmark αvβ8-blocking antibodies. Inhibition studies showed potent dose-dependent blockade by an anti-αvβ8 antibody. In contrast, pan-TGF-β neutralizing antibody displayed markedly weaker apparent potency, suggesting that targeting localized αvβ8-mediated activation is more effective than neutralizing released TGF-β in this assay context. The assay also enabled screening and ranking of anti-αvβ8 antibodies, identifying several high-potency clones, and detected αvβ8-mediated activation of a non-cleavable latent TGF-β1 mutant. This platform provides a sensitive, internally normalized, and scalable approach for mechanistic studies and therapeutic discovery targeting the αvβ8–TGF-β axis.

## Introduction

Transforming Growth Factor-β (TGF-β) is a multifunctional cytokine that regulates a wide range of cellular processes, including proliferation, differentiation, apoptosis, immune regulation, and extracellular matrix production^1,2^. Through these activities, TGF-β plays essential roles in embryonic development, tissue homeostasis, wound repair, and immune tolerance. Dysregulation of TGF-β signaling is implicated in numerous pathological conditions, including fibrosis^3^, inflammatory bowel disease^4^, cancer progression and metastasis^5^, neuroinflammation^6^, and autoimmune diseases^7,8^. Consequently, understanding the mechanisms governing TGF-β activation and signaling has become a major focus of therapeutic research.

Three closely related TGF-β isoforms, TGF-β1, TGF-β2, and TGF-β3, are encoded by distinct genes. They exhibit both overlapping and context-dependent biological functions^9^. TGF-β1 is the most abundant and extensively studied isoform and plays prominent roles in immune suppression, fibrosis, and tumor biology^10^. TGF-β2 exhibits more spatially restricted expression and is particularly important during embryonic development of the cardiovascular, skeletal, and nervous systems^11^. TGF-β3 plays critical roles in palatogenesis, lung development, and wound healing, where exogenous administration has been associated with reduced scarring in rat vocal fold mucosal injury^12^. Despite these functional distinctions, all three isoforms signal through the same receptor system and converge on canonical SMAD-dependent transcriptional pathways^13^. As TGF-β1 is the most abundant isoform and plays prominent roles in key disease areas such as immune suppression, fibrosis, and tumor biology, we prioritized the development of an assay specifically for latent TGF-β1 activation. The core dual-luciferase reporter system is nonetheless readily adaptable for studying the activation of TGF-β2 or TGF-β3 in future work.

A defining feature of TGF-β biology is that the cytokine is synthesized and secreted in a latent, inactive form^14^. Newly synthesized TGF-β is produced as a precursor. In this precursor, the mature TGF-β dimer is non-covalently associated with its latency-associated peptide (LAP)^14^. This complex is often associated with latent TGF-β binding proteins (LTBPs). This association forms a large latent complex that is deposited in the extracellular matrix (ECM)^15^. Latent sequestration provides a critical regulatory checkpoint, ensuring that TGF-β signaling is activated only within specific spatial and temporal contexts.

In addition to ECM sequestration, latent TGF-β can be presented on the cell surface by specialized milieu proteins. Glycoprotein A repetitions predominant (GARP, also known as LRRC32) anchors latent TGF-β1 on the surface of regulatory T cells and other cell types, enabling localized activation during cell–cell interactions^16^. Similarly, LRRC33 presents latent TGF-β1 on myeloid cells such as microglia and macrophages, contributing to immune regulation and tissue homeostasis^17^. These cell-surface presentation mechanisms are increasingly recognized as critical determinants of TGF-β bioavailability and function.

Activation of latent TGF-β is a tightly regulated and mechanistically diverse process^18^. Several activation mechanisms have been described. These include proteolytic cleavage of LAP by matrix metalloproteinases^19^ or plasmin, conformational changes induced by protein–protein interactions, and mechanically driven activation mediated by cellular forces. Among these mechanisms, integrin-mediated activation is particularly well characterized. Specific αv-containing integrins recognize an RGD motif within LAP and trigger localized TGF-β activation at the cell surface^20^.

Integrin αvβ8 is a major activator of latent TGF-β1^21^. It is highly expressed in specific cellular contexts, including neural, epithelial, and immune cells. In addition, it plays essential roles in central nervous system development, immune regulation, and tumor biology. Dysregulated αvβ8-mediated TGF-β activation has been implicated in cancer progression, neuroinflammatory disorders, and immune evasion, making αvβ8 an attractive therapeutic target^22^.

The hypothesis that αvβ8 activation relies solely on proteolytic cleavage has been challenged by previous studies utilizing the non-cleavable latent TGF-β1 mutant (R249A)^23^. This key mutation is known to prevent furin-mediated release of mature TGF-β1 from the latent complex, yet its activation by αvβ8 in other systems suggests that signaling can occur while the cytokine remains tethered. This observation highlights a critical mechanistic ambiguity regarding the role of cleavage in αvβ8-mediated activation.

Given the growing interest in targeting αvβ8-mediated TGF-β activation, there is a critical need for robust and physiologically relevant assays that directly measure this process. Foundational reporter assays using mink lung epithelial cells (MLECs)^24^ or HEK-Blue-TGFβ cells^25^ have been instrumental in advancing our understanding of TGF-β biology. However, these assays typically rely on the addition of exogenous TGF-β1 and the measurement of downstream signaling independently of the activation process. These approaches do not capture the unique biology of LTGF-β presentation and integrin-dependent activation. In addition, recent structural studies and assay platforms have employed immobilized recombinant integrin ectodomains to trigger activation^23^. This type of in vitro study without cell-based activation is a reductionist strategy that may not fully recapitulate physiological cell–cell interactions. Consequently, there remains a need for robust, internally normalized, cell-based assays capable of quantitatively measuring integrin-mediated TGF-β activation in a high-throughput format. Moreover, binding assays alone are insufficient to evaluate functional inhibition of αvβ8-mediated TGF-β activation, underscoring the importance of cell-based functional assays that faithfully recapitulate this complex activation mechanism. To address this gap, we developed a novel, fully cell-based co-culture assay that integrates cell-surface presentation of latent TGF-β1 with a ratiometric dual-luciferase reporter system to provide a direct, internally normalized, and physiologically relevant functional readout of integrin αvβ8-mediated activation.

In this study, we describe the development and validation of a novel cell-based reporter assay designed to quantitatively measure integrin αvβ8-mediated activation of latent TGF-β1. The assay employs a SMAD-responsive dual-luciferase reporter system combined with a co-culture model in which latent TGF-β1 is presented on the reporter cell surface by GARP. Activation of latent TGF-β1 by αvβ8 expressed on adjacent cells induces SMAD signaling and NanoLuc luciferase expression, while constitutive firefly luciferase activity provides an internal normalization control. This design enables sensitive, reproducible, and quantitative measurement of TGF-β1 activation in a physiologically relevant context.

We demonstrate that this assay reliably detects functional inhibition by benchmark αvβ8-blocking antibodies and it enables precise quantification of αvβ8-dependent activation of latent TGF-β1. Moreover, the assay is readily adaptable for high-throughput screening. This capability also allows for comparative evaluation of therapeutic antibodies and other modulators of the αvβ8–TGF-β pathway. Together, these features establish this assay as a versatile platform for mechanistic studies and drug discovery efforts targeting TGF-β activation.

## Materials and Methods

### Cell lines and culture conditions

NIH/3T3 (mouse embryonic fibroblast) and LN-229 (human glioblastoma) cells were obtained from the gCell repository (Genentech). NIH/3T3 cells were used to generate a stable SMAD-responsive TGF-β1 reporter cell population. LN-229 cells served as a source of endogenous integrin αvβ8 for co-culture activation assays. This cell line was specifically chosen for its confirmed high endogenous expression of αvβ8 (Figure 2B), which supports a physiologically relevant trans-activation model. Utilizing an endogenously expressing cell line, rather than a transfected one, ensures that the αvβ8 is correctly membrane-embedded and accurately recapitulates the cell–cell interactions required for activation, thereby expanding the utility of the co-culture design for mechanistic and drug discovery studies.

Cells were maintained in Dulbecco’s Modified Eagle Medium (DMEM, high glucose, 4.5 g/L) supplemented with 10% fetal bovine serum (FBS; Gibco), 2 mM L-glutamine, 100 U/mL penicillin, and 100 μg/mL streptomycin. Cells were cultured at 37°C in a humidified incubator with 5% CO₂.

Stable reporter cells were maintained under selection using Zeocin (500 μg/mL; InvivoGen) and/or Hygromycin B (200 μg/mL; InvivoGen) as indicated.

Following thawing, cells were maintained for at least one passage in growth medium without selection antibiotics before use in functional experiments.

### Assay medium

Unless otherwise indicated, functional assays were performed in assay medium consisting of DMEM (high glucose, 4.5 g/L) supplemented with 10% heat-inactivated FBS, 2 mM L-glutamine, 100 U/mL penicillin, and 100 μg/mL streptomycin.

## Construction of vectors

### TGF-β-responsive dual-luciferase reporter vector

A dual-luciferase reporter construct was generated to quantify canonical TGF-β/SMAD signaling, in which tandem SMAD-binding elements (SBE) are placed upstream of a minimal promoter (minP) to drive NanoLuc luciferase (Nluc) expression, enabling detection of SMAD pathway activation. A constitutively expressed bicistronic cassette driven by the human phosphoglycerate kinase (hPGK) promoter was incorporated into the vector backbone, in which the bleomycin resistance gene (bleoR) is followed by an internal ribosome entry site (IRES) and firefly luciferase (FLuc), enabling Zeocin selection and internal normalization. The resulting construct was designated SBE-NLucPest-FLuc-Zeo.

### Construction of GARP–latent TGF-β1 expression vectors

Two cell-surface latent TGF-β presentation constructs were generated by co-expressing human GARP (LRRC32) and full-length human latent TGF-β1 (LAP–TGF-β1) or latent TGF-β1 containing a point mutation (LTGF-β1 R249A) under the control of the CMV promoter. In these constructs, GARP was positioned upstream of a P2A peptide sequence followed by human latent TGF-β1 (LTGF-β1), enabling co-expression of both proteins from a single transcript. The R249A mutation results in expression of a form of TGF-β1 that cannot be released from the latent complex.

A hygromycin resistance gene driven by the SV40 promoter was included as the selection marker for stable mammalian cell expression in both constructs.

### Generation of stable 3T3 TGF-β reporter cells

TGF-β reporter cells were generated by stable transfection of NIH/3T3 cells with the SBE-NLucPest-FLuc-Zeo construct using Lipofectamine 3000 according to the manufacturer’s protocol. Forty-eight hours post-transfection, cells were subjected to Zeocin selection (500 μg/mL) for approximately 14 days.

Stable or transient transfection was subsequently performed with constructs expressing human GARP and LTGF-β1 (WT) or human GARP and LTGF-β1 (R249A). Stably transfected cells were maintained in the medium described in “Cell lines and culture conditions” for about 14 days to obtain a stable cell population, while transiently transfected cells were maintained in a similar medium without antibiotic selection.

### Antibodies and reagents

The following antibodies were used in this study: anti-human αvβ8 GNE Ab1 (mIgG2a) and GNE Ab2 (mIgG2a or mIgG2a, LALAPG); APC anti-human GARP (BioLegend, Cat# 352506); APC mouse IgG2b κ isotype control (BioLegend, Cat# 400320); PE anti-human LAP (BioLegend, Cat# 364404); PE mouse IgG1 κ isotype control (BioLegend, Cat# 400114); anti-αvβ8 GNE clone A (Rabbit IgG or mIgG2a.LALAPG); Alexa Fluor® 488 AffiniPure® F(ab’)₂ Fragment Goat Anti-Mouse IgG (H+L) (Jackson ImmunoResearch Laboratory Inc., Cat# 115-546-146). Anti-TGFβ pan inhibitor (GNE clone 1D11, hIgG4). The Nano-Glo® Dual-Luciferase Reporter Assay System (Promega, Cat# N1620) was used.

### Flow cytometry

Cell surface expression of GARP, LTGF-β1, and αvβ8 was analyzed by flow cytometry. Reporter cells were stained to assess co-expression of GARP and LTGF-β1. Cells were harvested and resuspended in the FACS wash buffer (PBS containing 1% BSA). For surface staining, cells were incubated with APC-conjugated anti-human GARP antibody and PE-conjugated anti-human LAP antibody, or the corresponding isotype control antibodies, for 30 min at 4 °C in the dark. Following incubation, cells were washed three times with 100 μL FACS wash buffer and resuspended in the same buffer for analysis.

Expression of αvβ8 was analyzed in LN-229 cells. Cells were harvested and resuspended in FACS wash buffer (PBS containing 1% BSA), and incubated with anti-αvβ8 antibody, Ab1, for 30 min at 4 °C. After washing three times with 100 μL FACS wash buffer, cells were incubated with Alexa Fluor® 488 AffiniPure® F(ab’)₂ fragment goat anti-mouse IgG (H+L) secondary antibody for 30 min at 4 °C in the dark. Cells were washed again and resuspended in FACS wash buffer prior to analysis.

Data were acquired using a CytoFLEX flow cytometer, with ≥10,000 events collected per sample, and analyzed using FlowJo software (BD Biosciences).

### αvβ8-mediated latent TGF-β1 activation co-culture assay

LN-229 effector cells were seeded at 40,000 cells per well in a 96-well white clear-bottom cell culture plate (Corning, Cat# 3610) in 50 μL assay medium and allowed to adhere for 1 h.

Stable 3T3 reporter cells expressing dual luciferase and GARP–LTGF-β1 were added at different numbers of cells per well in 30 μL assay medium. The final assay volume was 80 μL per well. Co-cultures were incubated for 18–20 h at 37°C in a humidified incubator with 5% CO₂.

To measure the quantity of NanoLuc luciferase and firefly luciferase, ONE-Glo™ EX reagent was pre-warmed to room temperature, and 80 μL per well was added to the cells. After an incubation of 20 minutes at room temperature with strong agitation, the firefly luminescence was measured with an EnSight multimode plate reader (PerkinElmer; Waltham, MA). Then 80 μL per well of NanoDLR™ Stop & Glo reagent was added to the plate. After an incubation of 20 minutes at room temperature with strong agitation, the NanoLuc luciferase luminescence was measured with an EnSight multimode plate reader. The NanoLuc luciferase signal was normalized to the firefly signal unless otherwise noted.

While initial assays for optimization were performed using a single NanoLuc luciferase readout, all rabbit antibody screening data reported herein utilized a dual-luciferase system. The inclusion of a constitutive firefly luciferase reporter provided an internal control for normalization, ensuring that the reported percent activation was independent of minor fluctuations in cell number and cellular variability.

### Antibody inhibition assays

Antibody inhibition assays were designed to quantify the functional inhibitory activity of therapeutic candidates against integrin αvβ8-mediated TGF-β1 activation. This enabled the precise screening and ranking of high-potency blocking antibodies.

Test antibodies were serially diluted in assay medium. LN-229 cells (42 μL; 40,000 cells per well) were pre-incubated with 8 μL of antibodies for 60 min at 37 °C in a humidified incubator with 5% CO₂, prior to the addition of 30 μL of reporter cells (20,000 cells per well). Co-cultures were then incubated for 18–20 h at 37 °C in a humidified incubator with 5% CO₂.

Dual-luciferase activity was measured as described above. Half-maximal inhibitory concentration (IC_50_) values were calculated from the titration curves using a Prism [Inhibitor] vs. response – Variable slope (four parameters) model (GraphPad Software, San Diego, CA), and the percentage TGF-β1 activation was calculated with the equation: % TGF-β1 activation = 100 x [(X - MIN)/(MAX - MIN)]. The MIN and MAX are the normalized NanoLuc luciferase signal from reporter cells alone and reporter cells co-cultured with LN-229 cells in test medium, respectively.

## Data analysis

All data were analyzed using GraphPad Prism (version 10.0; GraphPad Software). For the current study, experiments were performed with n = 1–2 technical replicate sets and N = 1–2 independent experiments, depending on the assay.

## Results

### Design of a co-culture cell-based reporter assay for integrin αvβ8–mediated activation of latent TGF-β1

To establish a cell-based platform for measuring integrin-mediated activation of latent TGF-β1, we generated stable NIH/3T3 reporter cells expressing a SMAD-responsive dual-luciferase cassette together with surface-presented GARP–LTGF-β1. In this system, activation of TGF-β receptors and downstream SMAD signaling induces NanoLuc luciferase expression, while constitutive firefly luciferase expression serves as an internal normalization control (Figure 1).

**Figure 1.**
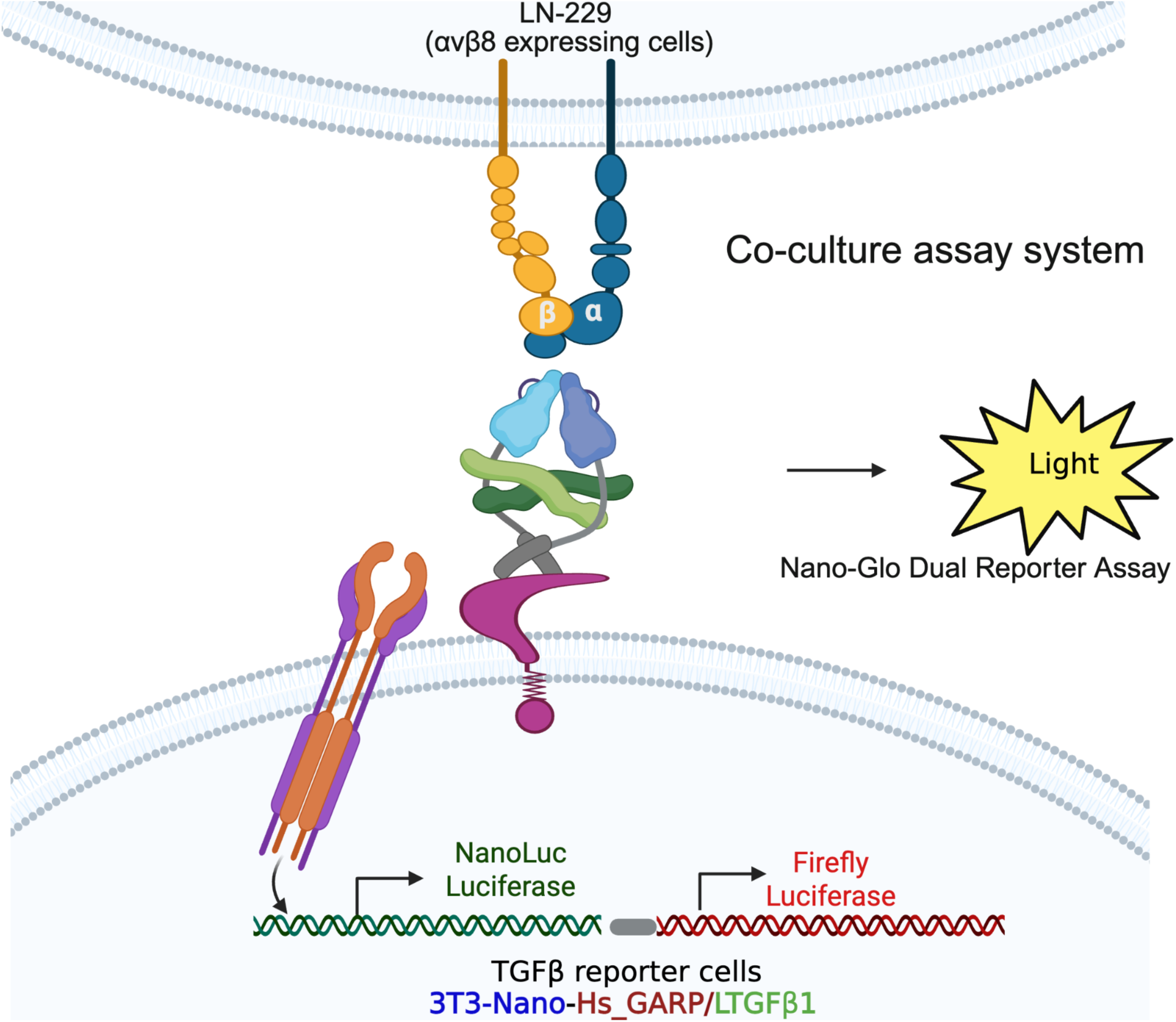
Schematic of the co-culture cell-based reporter assay for integrin αvβ8–mediated activation of latent TGF-β1. NIH/3T3 reporter cells express a SMAD-responsive NanoLuc luciferase reporter together with constitutive firefly luciferase for normalization and surface-presented GARP–latent TGF-β1. When co-cultured with LN-229 cells, which endogenously express integrin αvβ8, the integrin engages and activates GARP-presented LTGF-β1. Activated TGF-β1 subsequently stimulates TGF-β receptor signaling in the reporter cells, leading to SMAD pathway activation and induction of NanoLuc luciferase. Reporter activity is quantified as NanoLuc luciferase normalized to firefly luciferase. Created in BioRender. Zhang, J. (2026) https://BioRender.com/p4wkk60

To enable integrin-dependent activation of latent TGF-β1, the reporter cells were co-cultured with LN-229 cells, a glioblastoma cell line that endogenously expresses integrin αvβ8. In this co-culture configuration, αvβ8 mediates activation of latent TGF-β1 presented by GARP on the cell surface of reporter cells. This process leads to SMAD pathway activation and induction of the expression of NanoLuc luciferase reporter.

Together, the engineered reporter cells and αvβ8-positive LN-229 activator cells establish a robust cell-based assay platform used for all subsequent characterization and functional studies unless otherwise stated.

### Validation of cell-surface GARP–latent TGF-β1 and integrin αvβ8 expression

To confirm the key components of the co-culture assay, we first verified surface presentation of latent TGF-β1 on the engineered reporter cells. Flow cytometry using dual staining for human GARP and latent TGF-β1 demonstrated robust co-expression on NIH/3T3 TGF-β1–presenting reporter cells, with the majority of cells (97.5%) double positive relative to isotype controls (Figure 2A). These results confirm stable cell-surface presentation of the GARP–LTGF-β1 complex required for integrin-mediated activation.

**Figure 2.**
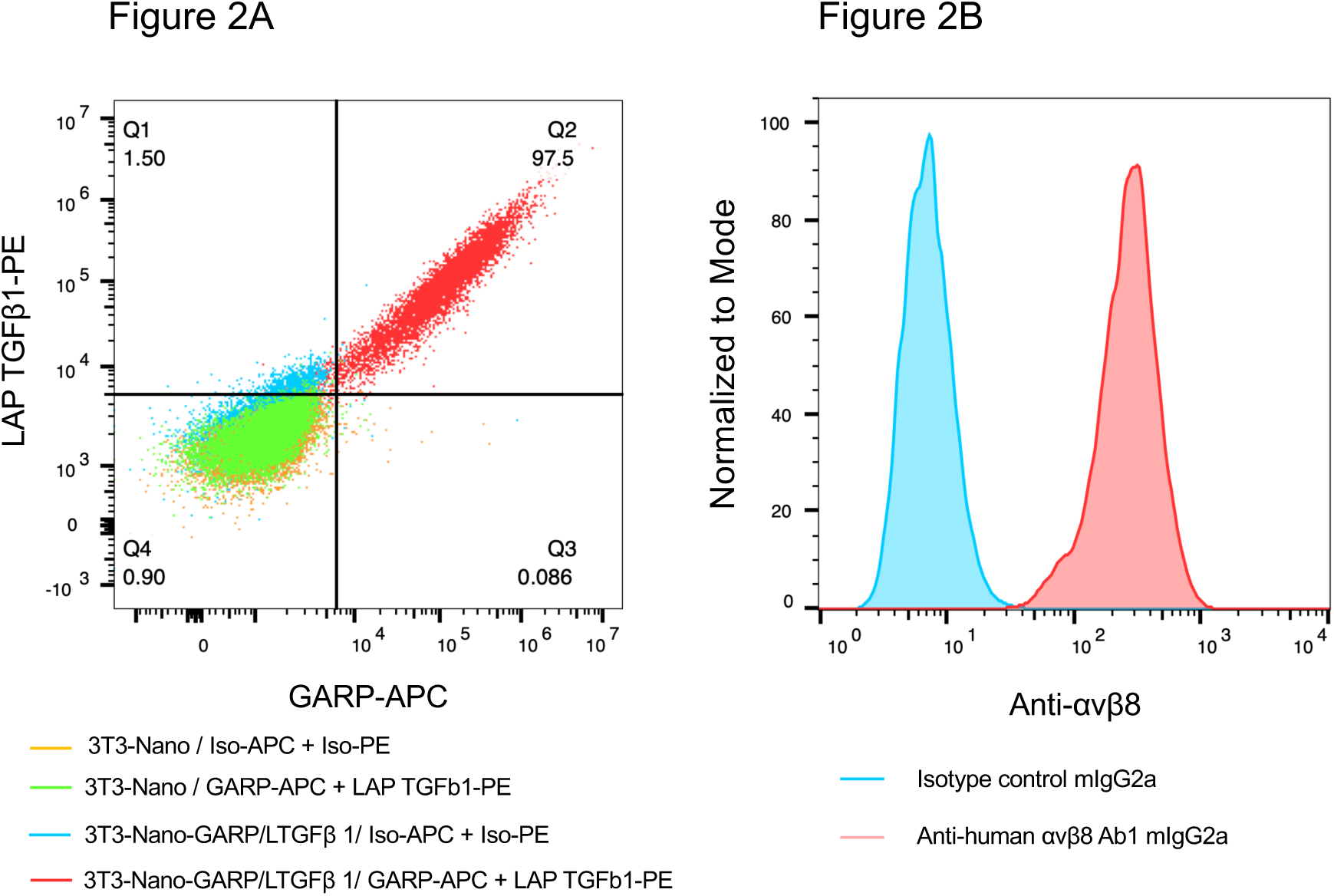
Flow cytometric validation of cell-surface GARP–latent TGF-β1 and integrin αvβ8 expression. **(A)** Dual-color flow cytometry analysis of engineered NIH/3T3 reporter cells stained for human GARP and latent TGF-β1. Representative plots show robust co-expression of GARP and LTGF-β1 on the cell surface compared with isotype controls. **(B)** Flow cytometry analysis of LN-229 effector cells stained with an antibody specific for integrin αvβ8 (Ab1), confirming endogenous surface expression relative to isotype control staining.

We next examined expression of the activator, integrin αvβ8, on the effector cells. Flow cytometry analysis showed that LN-229 cells endogenously express integrin αvβ8 on the cell surface (Figure 2B), supporting their use as activator cells capable of mediating LTGF-β1 activation in the co-culture reporter assay. Together, these results confirm that the biological components required for integrin αvβ8–mediated activation of latent TGF-β1 are present in the co-culture assay system.

### Optimization of αvβ8-mediated co-culture activation conditions

NanoLuc luciferase induction was dependent on both effector and reporter cell numbers. One of the critical requirements for this co-culture assay is the close contact between 3T3 reporter cells and effector LN-229 cells. Under these conditions, integrin αvβ8 expressed on the surface of LN-229 cells binds to LTGF-β1 presented by the reporter cells, enabling activation of the signaling pathway. To optimize cell density, particularly the number of reporter cells required to achieve an optimal signal-to-background ratio, the reporter cell density was systematically evaluated. LN-229 cells were seeded at a fixed density (40,000 cells per well) and co-cultured with TGF-β1 reporter cells at varying cell numbers (Figure 3). Background signal was determined as the signal from reporter cells only. Under these conditions, co-culture of 40,000 LN-229 cells with 20,000 reporter cells per well produced a signal-to-background ratio of 13.31, compared with ratios of 10.13 and 13.16 obtained with 40,000 LN-229 cells co-cultured with 40,000 or 10,000 reporter cells, respectively (Figure 3).

**Figure 3.**
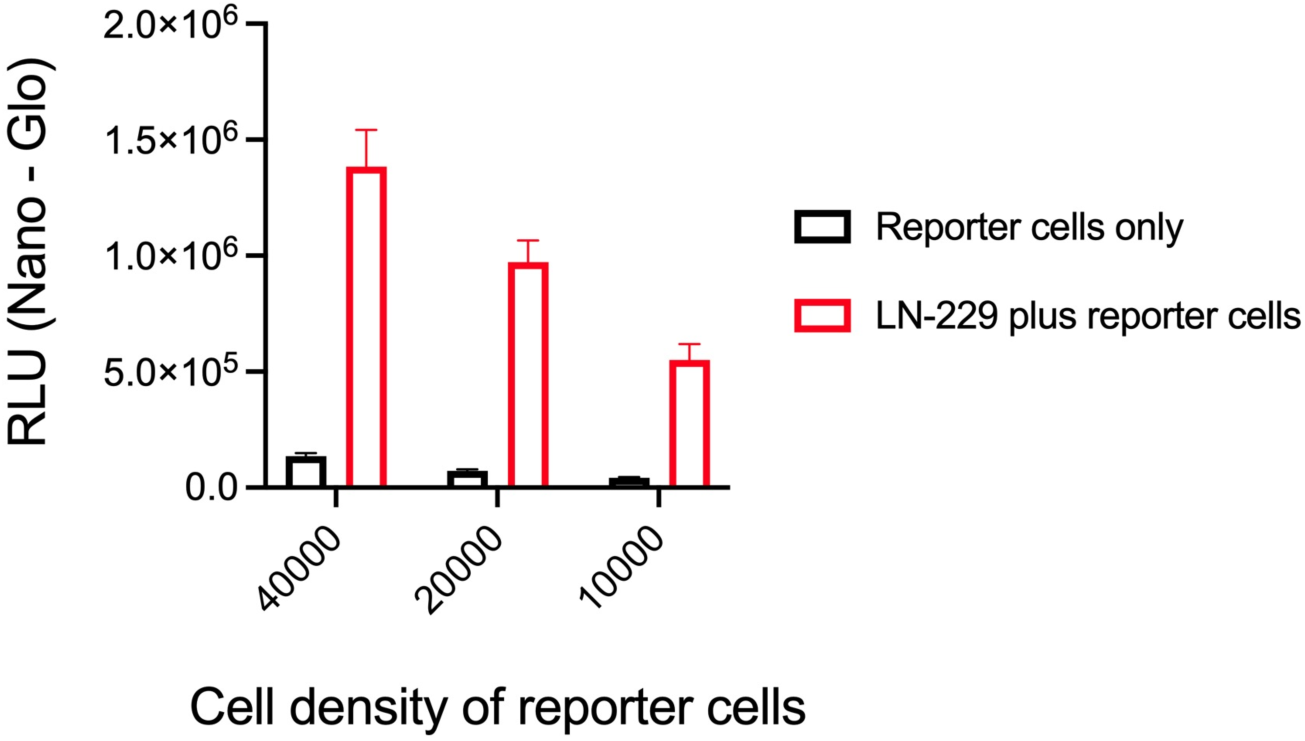
Optimization of reporter cell density for αvβ8-mediated LTGF-β1 activation. NIH/3T3 LTGF-β1 reporter cells were co-cultured with LN-229 effector cells at a fixed cell density. Reporter cell density was systematically varied to identify conditions yielding optimal NanoLuc luciferase signal and signal-to-background ratio. Data are shown as mean ± SD of 8 repeats from one experiment.

Based on these analyses, the condition of 40,000 LN-229 cells and 20,000 reporter cells per well was selected for subsequent experiments, as it provided a balance between assay robustness and practical throughput.

### Assay performance characteristics and αvβ8-dependent activation

To evaluate assay robustness and reproducibility, signal-to-background (S/B) ratios were determined across three independent assay runs. Robust activation was consistently observed in co-culture compared with reporter cells cultured alone. Mean S/B ratios were 8.4 ± 0.8, 10.2 ± 1.1, and 10.5 ± 1.2 across 3 independent experiments, with coefficients of variation (%CV) of 9.0%, 10.9%, and 11.2%, respectively (Figure 4).

**Figure 4.**
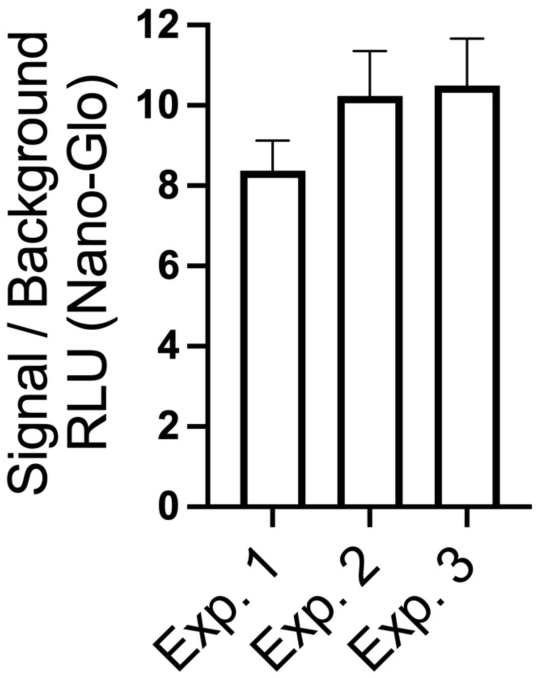
Assessment of assay robustness and reproducibility. Signal-to-background (S/B) ratios were determined across three independent assay runs under optimized co-culture conditions. The background represents reporter cells cultured alone in the absence of LN-229 cells. Bars represent mean ± SD (n = 4, 4, and 6 technical replicates for Assays 1, 2, and 3, respectively).

In this analysis, the signal represents NanoLuc activity measured under co-culture conditions, whereas the background represents reporter cells cultured alone in the absence of LN-229 cells. Together, these results demonstrate that the assay is robust and reproducible and that NanoLuc luciferase induction reflects αvβ8-dependent activation.

To further validate the assay for inhibitor screening and confirm integrin αvβ8–mediated activation, the anti-αvβ8 inhibitory antibody Ab2 and the pan–TGF-β neutralizing antibody 1D11 were evaluated in the co-culture system. The αvβ8-blocking antibody Ab2 inhibited reporter activation in a dose-dependent manner, yielding an IC₅₀ of 0.88 nM (Figure 5). In contrast, the pan–TGF-β neutralizing antibody 1D11 (known to neutralize all three active TGF-β isoforms) exhibited substantially weaker apparent potency, with an IC₅₀ of approximately 99.37 nM. These findings demonstrate that activation in this system is strictly integrin αvβ8–dependent. The marked difference in apparent potency between the αvβ8 blocker (Ab2) and the pan-TGF-β neutralizing antibody (1D11) supports the assay’s suitability for evaluating anti-αvβ8 therapeutic antibodies and highlights that the assay can discriminate among inhibitors with distinct potencies.

**Figure 5.**
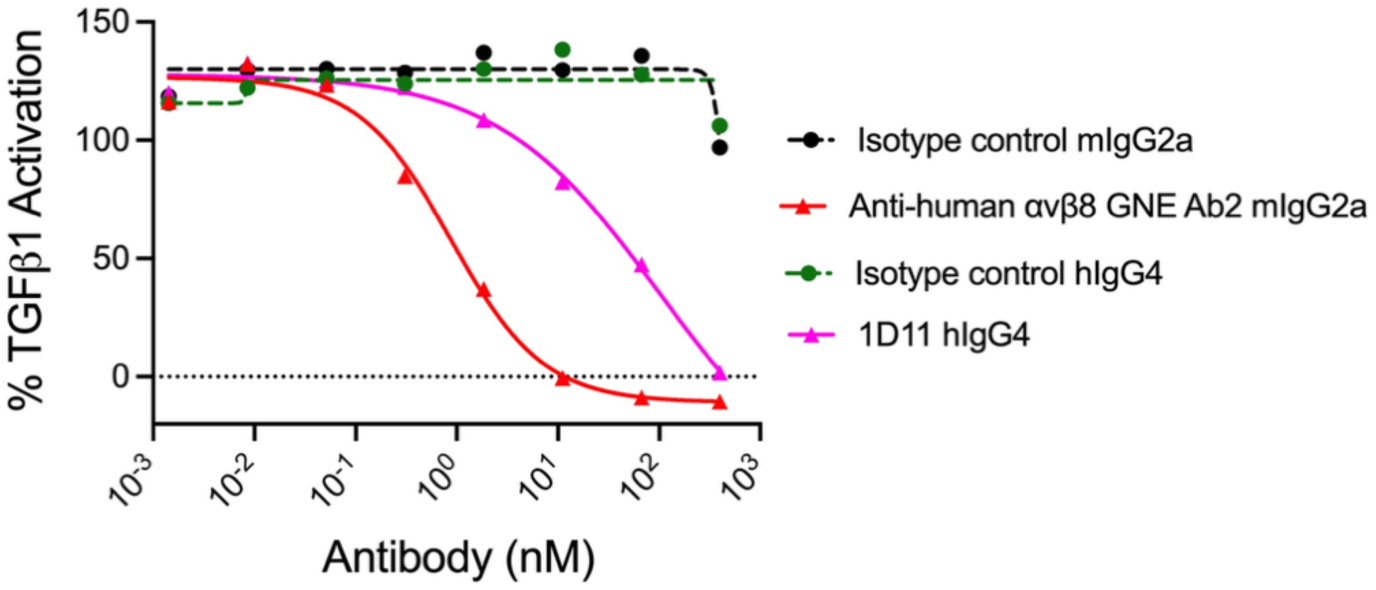
Antibody inhibition of αvβ8-mediated LTGF-β1 activation. Dose–response inhibition curves for anti-human αvβ8 antibody, Ab2, and 1D11 measured using the co-culture reporter assay. NanoLuc activity served as the readout of TGF-β signaling. IC₅₀ values were calculated by nonlinear regression (four-parameter logistic model).

To demonstrate the utility of the assay for antibody discovery, a panel of anti-αvβ8 antibodies was evaluated in a primary screening format at a single concentration of 10 μg/mL (66.6 nM). The isotype control showed 115.5% TGF-β1 activation, whereas the positive control antibody Ab2 showed −10.3% activation. The addition of the anti-αvβ8 antibody clones into the co-culture system modulated the integrin-mediated TGF-β1 activation signal, resulting in a wide range of calculated activities (from −16.7% to 133.9% activation, Figure 6A). The lower percentage values signify greater functional inhibition of the αvβ8 activation pathway. Antibodies producing less than 25% LTGF-β1 activation under these conditions were selected as hits. These hits advanced to confirmatory dose–response analysis for IC₅₀ determination and clone ranking. In the dose–response assays, antibodies were tested starting at 66.6 nM with an eight-point 1:6 serial dilution. Representative data are shown in Figure 6B.

**Figure 6.**
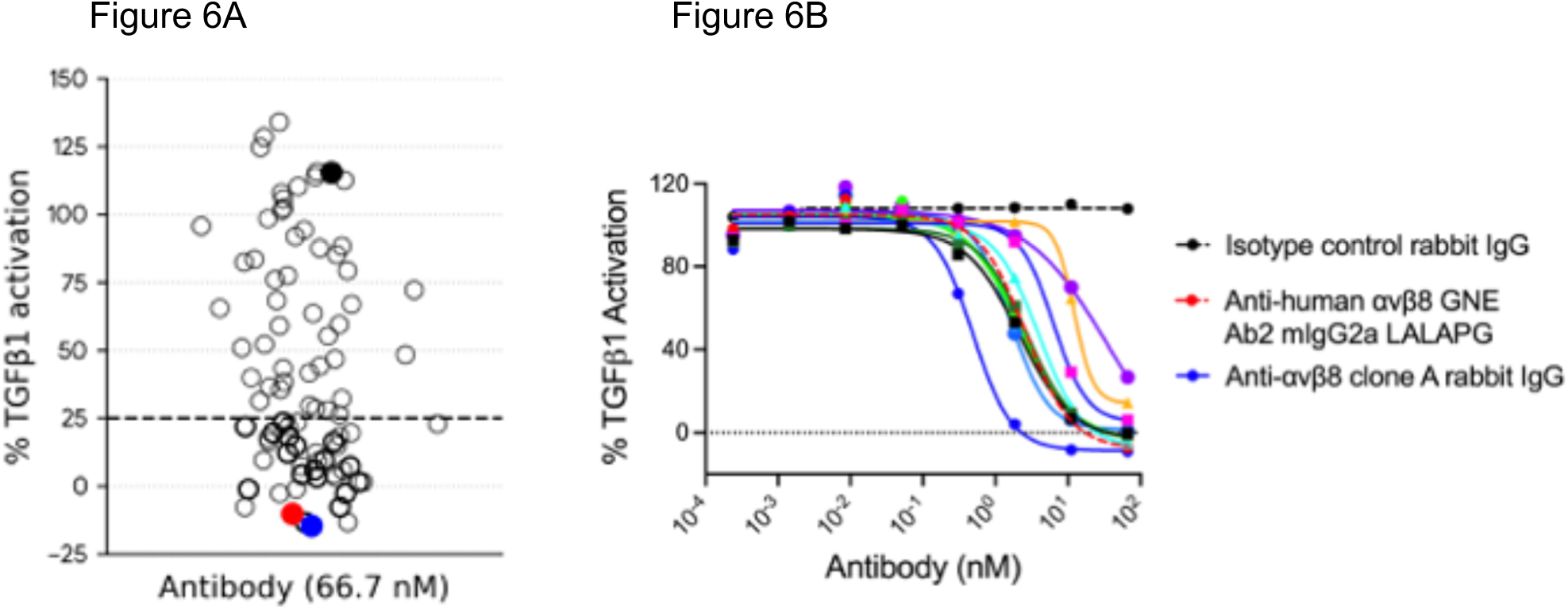
Application of the co-culture reporter assay for screening anti-αvβ8 rabbit antibodies. **(A)** Primary screening of a panel of anti-αvβ8 rabbit antibodies at a single concentration (10 µg/mL, 66.6 nM) using the co-culture reporter assay. Isotype control (black) and the anti-αvβ8-blocking antibody Ab2 (red) were included as negative and positive controls, respectively. Antibodies producing reduced LTGF-β1 activation were selected as hits for further evaluation. **(B)** Representative dose–response inhibition curves of selected anti-αvβ8 rabbit antibodies evaluated in the co-culture reporter assay. Antibodies were tested starting at 66.6 nM using an eight-point 1:6 serial dilution, and IC₅₀ values were determined by nonlinear regression using a four-parameter logistic model. Negative and positive controls are indicated by dashed lines.

The IC₅₀ values of the positive clones ranged from 0.49 to 33.95 nM, with some values estimated due to incomplete curve fitting. Clone A showed the strongest potency (IC₅₀ = 0.49 nM). These results demonstrate the utility of this platform for screening and identifying high-potency anti-αvβ8 antibody candidates for further development.

### Evaluation of integrin-mediated activation using a non-cleavable LTGF-β1 mutant

To further demonstrate the platform’s capability to evaluate mechanisms of integrin-mediated latent TGF-β1 activation, we examined a mutant LTGF-β1 construct carrying the R249A substitution in the latency-associated peptide (LAP)^14^. This mutation has been reported to prevent furin-mediated cleavage of mature TGF-β1 from the latent complex. Structural and functional analyses by Campbell and colleagues further suggested that αvβ8-mediated activation of latent TGF-β can occur without release and diffusion of mature TGF-β, supporting a model in which signaling occurs while TGF-β remains associated with the latent complex^23^.

To evaluate this mechanism within our assay system, engineered reporter cells expressing GARP-associated LTGF-β1 (R249A) were generated and tested using the co-culture reporter assay described above. When co-cultured with LN-229 cells expressing integrin αvβ8, reporter activation was readily detected, indicating that the assay can measure integrin-dependent activation of the non-cleavable latent TGF-β1 complex. Under these conditions, the αvβ8-blocking antibodies, clone A and GNE Ab2, strongly inhibited LTGF-β1 activation, with IC₅₀ values of 0.11 nM and 0.24 nM, respectively (Figure 7A). In contrast, the TGF-β pan inhibitor 1D11 exhibited substantially weaker inhibition than the αvβ8-blocking antibody GNE Ab2, with an IC₅₀ of 23.29 nM versus 0.26 nM for Ab2 (Figure 7B). These results demonstrate that the assay platform can also detect integrin αvβ8–mediated activation of the non-cleavable LTGF-β1 (R249A) complex. In this system, αvβ8-blocking antibodies inhibited reporter activation with substantially greater potency than the pan–TGF-β inhibitor 1D11, consistent with activation occurring in an integrin-dependent manner within the latent complex.

**Figure 7.**
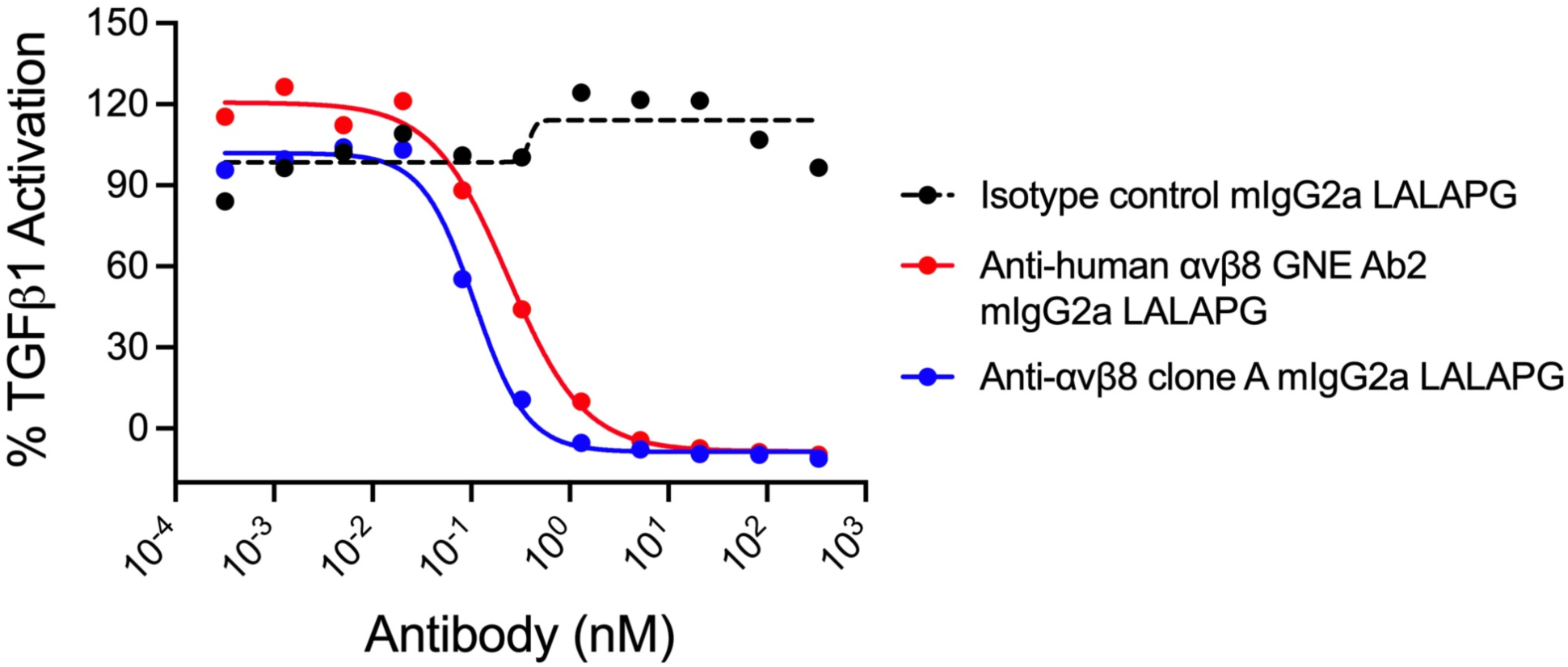
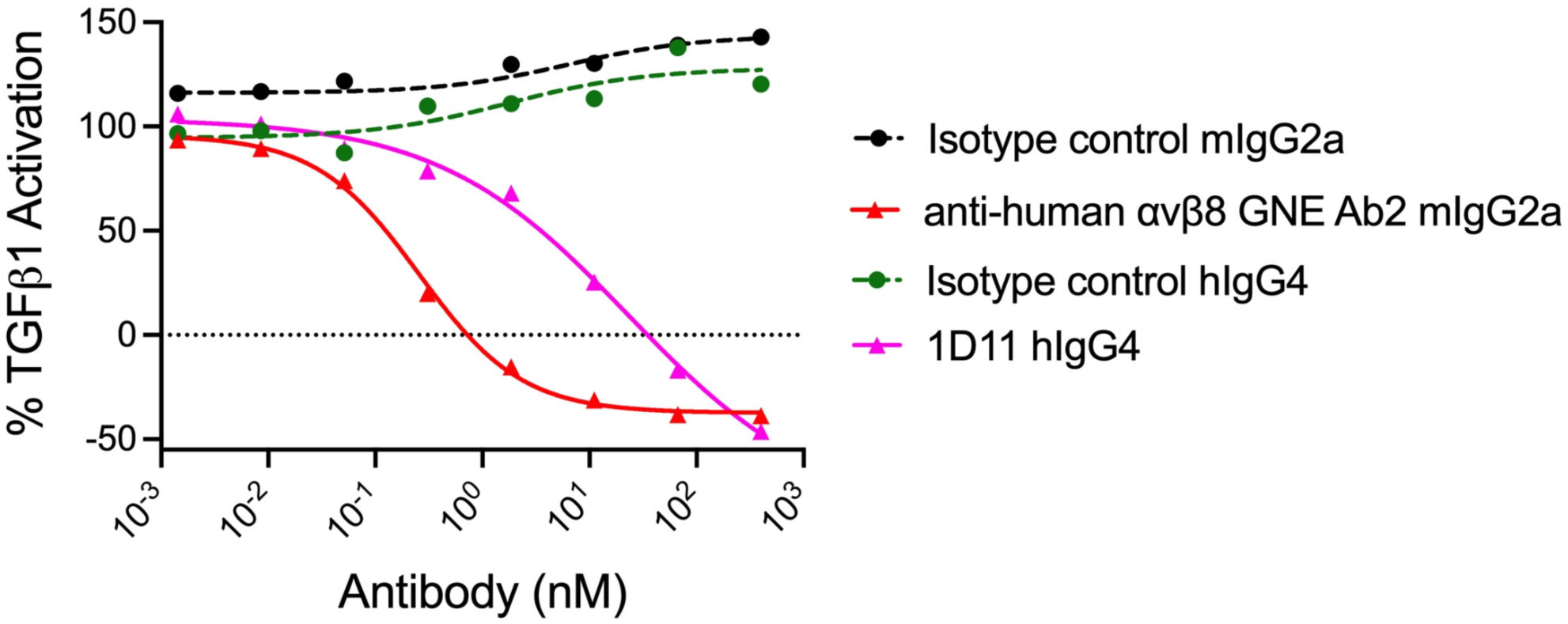
Evaluation of integrin αvβ8–mediated activation using a non-cleavable LTGF-β1 (R249A) mutant. Reporter cells transiently expressing GARP-associated LTGF-β1 (R249A) were co-cultured with LN-229 cells expressing integrin αvβ8. Antibodies were tested using an 11-point 1:4 serial dilution. IC₅₀ values were determined by nonlinear regression using a four-parameter logistic model. **(A)** Inhibitory activities of αvβ8-blocking antibodies clone A and GNE Ab2. **(B)** Comparison of inhibitory activities between the αvβ8-blocking antibody GNE Ab2 and the TGF-β pan inhibitor 1D11. The dose–response curves were generated using NanoLuc luciferase signals without normalization to firefly luciferase activity.

## Discussion

In this study, we developed and validated a co-culture cell-based reporter platform that enables quantitative measurement of integrin αvβ8–mediated activation of latent TGF-β1. By combining surface presentation of GARP-associated latent TGF-β1 with a SMAD-responsive NanoLuc luciferase reporter system, this platform directly links localized activation of latent TGF-β1 to a sensitive transcriptional readout. This design allows functional interrogation of integrin-dependent TGF-β1 activation in a physiologically relevant context and provides a robust assay suitable for mechanistic studies and therapeutic screening.

The engineered reporter cells exhibited stable co-expression of GARP and latent TGF-β1 and retained full responsiveness to canonical TGF-β signaling. Importantly, surface presentation of latent TGF-β1 did not impair downstream signaling competence, as demonstrated by robust SMAD-responsive reporter induction. These findings confirm that the reporter system captures authentic TGF-β1 signaling while enabling controlled investigation of the upstream integrin-dependent activation mechanisms.

The co-culture configuration recapitulates a key biological pathway in which αvβ8-expressing cells activate latent TGF-β1 presented by neighboring cells. In this system, integrin αvβ8 expressed on LN-229 effector cells interacts with GARP-associated latent TGF-β1 on reporter cells, triggering activation and downstream SMAD signaling. The strong dependence on functional GARP–LTGF-β1 presentation and the requirement for cell-cell interaction highlight the localized nature of TGF-β1 activation and reflect mechanisms believed to operate in physiological settings such as immune regulation^26^ and tumor microenvironments^27^.

Optimizing co-culture parameters improved assay robustness, sensitivity, and reproducibility, yielding strong signal-to-background ratios with low variability across independent runs. The ratiometric dual-luciferase design significantly improves assay precision relative to traditional single-reporter systems such as the mink lung epithelial cell (MLEC/TMLC) assays originally described by Abe et al.^24^ Whereas single-reporter systems require parallel cytotoxicity assays or external normalization to account for well-to-well variability in viability and cell number, the constitutive firefly luciferase provides internal normalization, improving measurement precision and supporting quantitative screening applications.

The assay also demonstrated clear dependence on integrin αvβ8 activity. Benchmark αvβ8-blocking antibodies inhibited reporter activation in a dose-dependent manner with subnanomolar potency, whereas antibodies targeting active TGF-β showed substantially weaker inhibition. Unlike binding-based assays, this platform directly measures biologically relevant TGF-β activation, enabling discrimination between antibodies that bind αvβ8 and those that effectively inhibit its functional activity. This distinction is particularly critical for drug discovery efforts targeting the TGF-β pathway, where binding affinity alone does not reliably predict functional blockade. The differential potency observed between the αvβ8 blocking antibody (Ab2) and the pan–TGF-β neutralizing antibody (1D11) highlights the utility of this αvβ8-dependent assay platform for functional evaluation of therapeutic antibodies. The significantly higher potency of the integrin-blocking antibody suggests that targeting localized αvβ8-mediated activation is more effective than neutralizing released TGF-β in this assay context. Although integrin αvβ6 is also a major activator of latent TGF-β1, the anti-αvβ8 leads used in this study were selected based on extensive binding specificity screens. These assays confirmed no detectable binding to human αvβ1, αvβ3, and αvβ5 at 100 nM, and no binding to human αvβ6 at 300 nM. This robust binding selectivity supports the use of these antibodies in specifically demonstrating αvβ8-dependent activation in our co-culture system.

Compared with existing TGF-β reporter systems, this platform provides several advantages. This assay advances physiological relevance by employing a fully cell-based co-culture design rather than immobilized protein substrates. While previous mechanistic studies, including those by Campbell et al.^23^, which utilized TMLC reporter cells immobilized on purified integrin αvβ8 ectodomains, have provided valuable structural insights, such reductionist systems lack the membrane context and cell-cell interactions present in vivo. Our approach uses LN-229 cells to present endogenous, membrane-embedded αvβ8 to reporter cells. This trans-activation model more accurately reflects the cell–cell interactions characteristic of immune and tumor microenvironments and recapitulates physiological activation mechanisms.

Evaluation of the non-cleavable LTGF-β1 R249A mutant further demonstrates the mechanistic utility of the system. Despite preventing conventional proteolytic release of mature TGF-β^23^, this mutant still supported αvβ8-dependent reporter activation. These findings are consistent with models suggesting that integrin-mediated activation can occur without complete release of mature TGF-β from the latent complex, allowing signaling while the cytokine remains associated with its latency components.

This controlled design provides a reproducible, scalable, and adaptable framework for both mechanistic studies and screening efforts, although several limitations should be considered. The assay relies on engineered cell lines expressing GARP-associated latent TGF-β1, which may not fully reflect endogenous cytokine presentation levels. In addition, LN-229 cells serve as the source of integrin αvβ8 and therefore may not capture the diversity of cellular contexts in which integrin-mediated activation occurs in vivo. Incorporating additional integrin αvβ8-expressing cell types may further expand the physiological relevance of the system. Finally, future adaptations may incorporate alternative mechanisms for presenting latent TGF-β, additional integrin subtypes, or disease-relevant cellular contexts to further expand the platform’s utility.

## Conclusion

In this study, we developed and validated a robust cell-based assay that enables quantitative measurement of integrin αvβ8–mediated activation of latent TGF-β1. By combining physiologically relevant presentation of latent TGF-β1 with a sensitive SMAD-responsive dual-luciferase reporter system, the assay provides a direct functional readout of localized TGF-β1 activation. The platform demonstrates strong performance characteristics, including a wide dynamic range, high reproducibility, and suitability for antibody screening and lead optimization.

Collectively, this assay fills a critical gap in existing TGF-β1 activation technologies and offers a versatile tool for mechanistic investigation and therapeutic development targeting the αvβ8–TGF-β1 axis. With its scalability and adaptability, the platform has the potential to accelerate the discovery of novel therapeutic strategies for diseases driven by dysregulated TGF-β1 activation.

## Acknowledgements

The authors thank Isabelle Lehoux and the BioMolecular Research Molecular Biology group for their expertise and assistance in DNA construct generation. Also, thank you to the following for help with the critical review of this manuscript: Arturo Orjalo, Yih-Wen Chen, Jerry Wang, and Mercedesz Balazs.

## Notes

### Competing Interest Statement

All the authors are current or previous employees and shareholders of Roche/Genentech

